# Within-host viral evolution varies with T cell receptor repertoire diversity independently of viral mutation rates

**DOI:** 10.1101/2025.05.19.654678

**Authors:** Hassan Jamaleddine, Anmar Khadra, Judith N. Mandl

## Abstract

The specific T cell receptors (TCRs) present in an individual determine which antigens can be recognized, and hence a repertoire comprised of a vast number of unique TCRs ensures an effective T cell response can be mounted against a broad range of potential pathogens encountered. To evade the T cell response, pathogens, and viruses in particular, can accrue epitope escape mutations, establishing a host-pathogen ‘tug-of-war’ between T cell-mediated control and viral evolution. Indeed, the evolutionary significance of TCR diversification in mammals is underscored by the sophisticated molecular machinery – spanning from somatic V(D)J recombination through non-templated nucleotide additions – that establish the TCR repertoire. Yet, whether greater oligoclonality among responding T cells, as is the case for elderly or immunocompromised individuals, might therefore accelerate pathogen immune escape remains unclear. Here, we constructed a computational model of T cell-mediated viral evolution to investigate how the number of unique virus-specific T cells clones impacts the balance between curtailing viral load through cross-reactivity to mutant strains on the one hand, versus promoting the generation of escape variants on the other. The model recapitulated prior experimental findings, such as T cell-dependent emergence of new viral mutants and the progressive decrease in the number of T cell clones present in the response over time. Our model simulations suggested that oligoclonality in the virus-specific T cell response can, if coupled with a resulting increase in viral replication, maximize viral immune escape despite constant mutation rates revealing a mechanism for how viruses with comparable mutation rates can nonetheless exhibit dramatically different adaption rates within hosts. Taken together, the model results suggest that variations in TCR repertoire diversity, and consequently in the number of pathogen-specific T cell clones within an individual, can have direct, non-linear impacts on viral immune escape.

## Introduction

T cells are key mediators of anti-viral immune responses, able to recognize and respond to viral pathogens in an antigen-specific manner through the interaction between their T cell receptors (TCRs) and virus-derived peptides presented by major histocompatibility complex molecules (pMHC). The T cell population within an individual is composed of a highly diverse set of TCRs that are generated by V(D)J recombination of the TCRα and β chain genes during T cell development in the thymus (1-3). The collection of unique CD8^+^ TCRs that recognize any given viral pMHC epitope is typically in the range of around 10^2^-10^3^ T cell precursors (4-6), with individual TCR sequences differing in their frequencies within and across individuals (7) and in their propensity for cross-reacting to different pMHC epitopes (6, 8). In all, the number of unique sequences that make up the total TCR repertoire of an individual has been estimated to be around 10-100 million in humans (9-11). Given the breadth of possible viral pathogens an individual might encounter, this diversity in TCR repertoire composition is presumably necessary to ensure a response can be mounted irrespective of the virus in question. Importantly, however, what constitutes a sufficiently diverse TCR repertoire remains poorly defined.

To counter TCR-driven T cell responses, viruses have evolved the ability to escape T cell immunity by acquiring genome mutations that result in altered epitope sequences and reduced presentation by MHC and/or reduced binding by specific TCRs (12-14). In the case of chronic viruses such as human immunodeficiency virus (HIV) and hepatitis C virus, rates of within-host viral evolution and cytotoxic T cell (CTL) escape are substantial, outpacing the development of an effective immune response, leading to persistent infection in the host and posing a significant barrier in the development of effective vaccines (15-19). Interestingly, HIV-1 has comparable mutation rates to those of other RNA viruses when measured in number of substitutions per nucleotide per infection cycle, yet demonstrates far greater rates of within-host evolution within TCR epitopes than, for instance, lymphocytic choriomeningitis virus (LCMV) in mice, where TCR epitope-restricted mutations are observed much less consistently, and at lower frequencies (20, 21). Thus, questions remain as to how TCR-driven selection pressure interacts with pathogen intrinsic dynamics to maximize the propensity for viral mutants to arise, irrespective of the rate of mutation, and whether the number of distinct virus-specific TCR clones resulting from varying the total diversity of the TCR repertoire might play a decisive role.

Understanding how the generation of viral T cell epitope mutants varies as a function of the number of responsive virus-specific TCRs in the TCR repertoire is imperative given the large, multi-factorial variability in the total repertoire diversity and, consequently, in the potential number of pathogen-specific TCRs among individuals. For example, thymic involution with age is known to lead to decreased T cell production and a contraction in TCR repertoire diversity with time (10, 22). Additionally, TCR repertoire diversity declines following various forms of immunological injury such as HIV-induced CD4^+^ T cell depletion or hematopoietic stem cell reconstitution (23-25). Given the potential for this variability to differentially drive within-host viral adaptation, it becomes important to investigate how varying the number of unique responding TCR clonotypes itself, given different levels of total TCR repertoire diversity, can impact the generation of viral escape mutants.

Interestingly, some studies have described a correlation between the number of unique TCRs of responding T cells specific for viral pMHC epitopes and T cell immune escape. For example, TCR sequencing of hepatitis C virus (HCV)-specific T cells in chimpanzees and of simian immunodeficiency virus (SIV)-specific T cells in rhesus macaques were generally associated with more oligoclonal TCR responses and with higher rates of CTL escape during chronic infection (26, 27). Similarly in humans, HIV quasispecies diversity in chronically infected individuals was shown to vary with HIV-specific T cell diversity, with greater TCRβ chain diversity inversely correlating with HIV evolution within-host (28). However, whether the TCR diversity of virus-specific T cells could *causatively* modulate the rates of within-host viral evolution and immune escape, and the manner in which this effect might arise from the dynamics of immune-viral interactions, has not been established.

In this work, we develop a computational model to theoretically explore the relationship between within-host evolution of a mutating virus and the TCR diversity of the responding T cell population. We show that varying the number of unique virus-specific clones can drive pathogen loads to regimes that maximize the propensity of the virus to generate escape mutants. Thus, our work shows that TCR diversity can critically impact the level of within-host viral evolution independent of viral mutation rate.

## Results

### Computational model simulates within-host viral evolution as a function of TCR repertoire diversity

To study the effect of the diversity of reactive TCRs on viral immune escape, we developed a computational model of viral replication and mutation in the presence of antigen-specific T cells with a varying number of unique TCR clonotypes, *N* (**Fig. 1A**, see Methods for detailed model description). T cell dynamics in the model includes terms for: constant input from the thymus, virus-induced proliferation, natural turnover of T cells (death rate), and intra- and inter-clonal competition for antigen and resources (with the former competition rate assumed to be greater than the latter). To incorporate within-host viral evolution, the initial parental strain can give rise to mutant daughter strains throughout the infection (**Fig. 1B**), with a mutation rate that is proportional to the abundance of the parental strain (i.e., strains with high pathogen loads are more likely to generate mutant progeny than strains with lower pathogen loads). Here, a mutant refers specifically to a new viral strain with an altered pMHC epitope presented to virus-specific TCRs. To every pair of T cell clones and viral strains, we assigned a value representing the specificity of a given clone for a given viral strain. Upon mutation, the specificity of any T cell clone for the daughter strain was assumed to be a small deviation in magnitude from its specificity for the parental strain (**Fig. 1C**); this assumption is supported by previous observations that TCR signal strength in response to a given pMHC epitope generally correlates with signal strength to epitopes with single amino acid mutations (6, 29).

**Figure 1.**
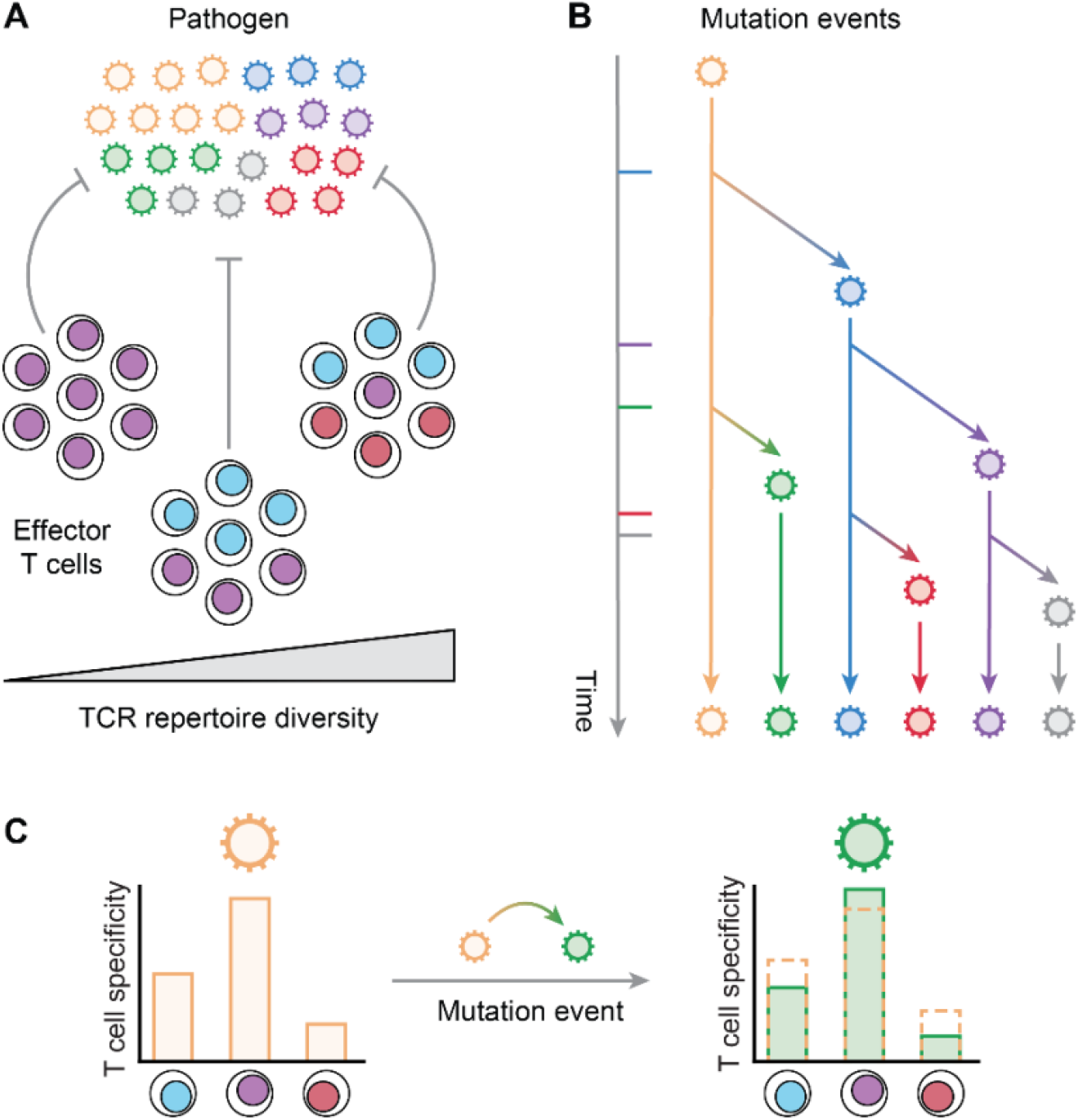
Computational population model of effector T cell diversity in response to an evolving viral pathogen. **(A)** Schematic illustrating effector T cells of varying TCR repertoire diversity responding to different variants of a replicating virus. **(B)** The virus is allowed to mutate within the model; the likelihood of a mutation event is proportional to the abundance of the parent strain. **(C)** Each T cell clone is assigned a value equivalent to the specificity of that clone for a pathogen strain. Upon mutation, the specificity of a T cell clone for a mutant strain is perturbed relative to its specificity for the parent strain.

By simulating the model with one initial strain of the virus and *N* = 100 unique TCR clonotypes, we generated time series showing different strains of the virus emerging in succession (**Fig. 2A**) and the resulting T cell response across all clonotypes (**Fig. 2B**). To measure the diversity of the responding T cell population over time, we computed the number of clonotypes that represented the majority (95%) of the total T cell population at early and late infection time points. We found that clonotype diversity contracts at later time points relative to peak infection (**Fig. 2C**). This observation in our model is consistent with measurements made in early vs. late stages of infection of mice with a chronic strain of LCMV (clone 13) (30); thus, the computational model was able to recapitulate TCR clonotype dynamics in the context of chronic infection and viral evolution under T cell pressure.

**Figure 2.**
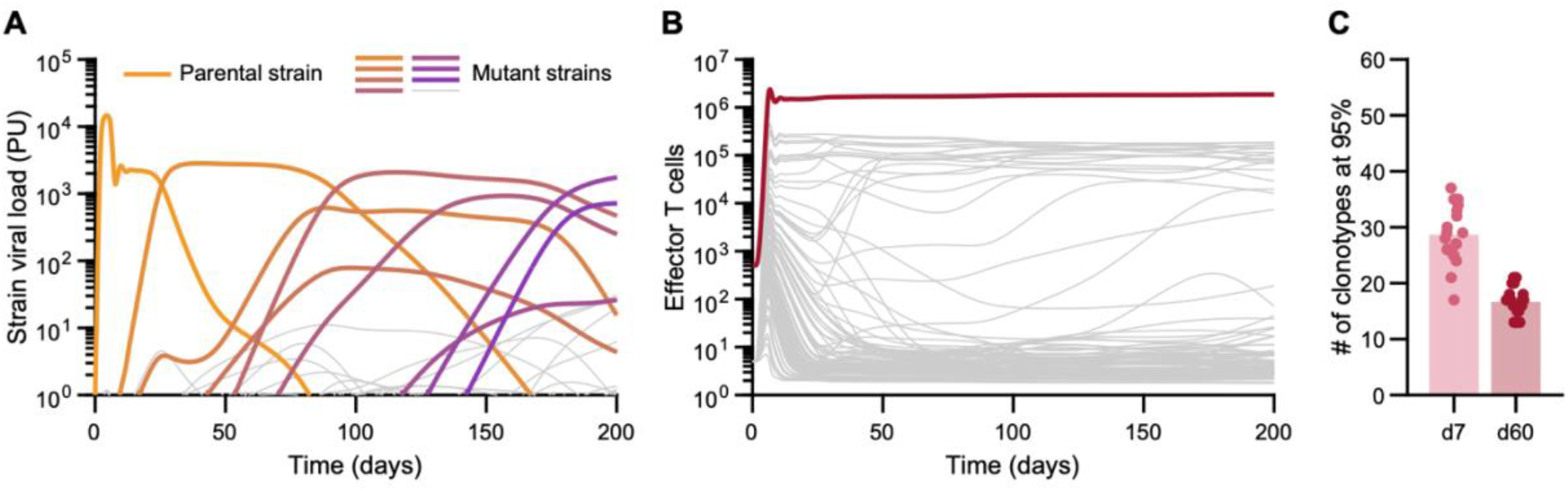
Model simulations of the T cell response to mutating pathogen captures contraction of clonotype diversity over time. **(A)** Pathogen load for each individual strain obtained from a single realization simulated by the model, starting from a single parental strain (Strain 1). Dominant strains (colored curves) are distinguished from those that have low abundance (gray curves) constituting < 2% of the pathogen pool at any given time. **(B)** Total number of effector T cells over time (red curve), comprised of 100 individual T cell clonotypes (gray curves) obtained from one single realization simulated by the model. **(C)** Number of T cell clonotypes that make up 95% of the responding T cell population at early vs. late stages of a chronic infection, computed from 50 independent realizations of model simulations.

### The presence of a T cell response is required for dominant viral mutants to replace the parental strain

Using our model framework, we next investigated how mutational burden (i.e., the number and abundance of emerging viral variants) differs in the presence vs. absence of T cell selective pressure in our simulations. Previous work showed that the frequency of LCMV-Cl13 viral TCR epitope mutants 39 to 75 days post-infection in RAG2^−/–^ mice (lacking both functional T and B cells) are negligible when compared to those in WT mice (31). To test whether our model is consistent with these data, we first simulated 200 independent realizations of the model in the absence of T cells or with *N* = 100 unique T cell clonotypes, starting with a single parental strain, and quantified the cumulative number of strains of the virus that emerged over the course of each simulation (**Fig. 3A, B**). In contrast to the data from LCMV-Cl13 infections in RAG2^−/–^ mice, we observed a greater absolute number of mutants over the course of an infection when T cells were absent in the model (**Fig. 3B**). In fact, the number of total strains mirrored the simulated pathogen loads, with higher pathogen replication obtained in the absence than in the presence of T cells (**Fig. S1A**). To experimentally test whether a virus replicating in the absence of T cell control results in higher viral loads, we infected WT and TCRβ KO mice with chronic LCMV-Cl13 and quantified viral titre via quantitative PCR 21 days post-infection. We found that, indeed, TCRβ KO mice possessed higher LCMV-Cl13 glycoprotein (GP) RNA levels relative to WT mice (**Fig. S1B, C**). This direct relationship between pathogen load and the total strain count in the model was not surprising given the model design, wherein the rate of mutation was assumed to scale linearly with pathogen abundance in the model.

**Figure 3.**
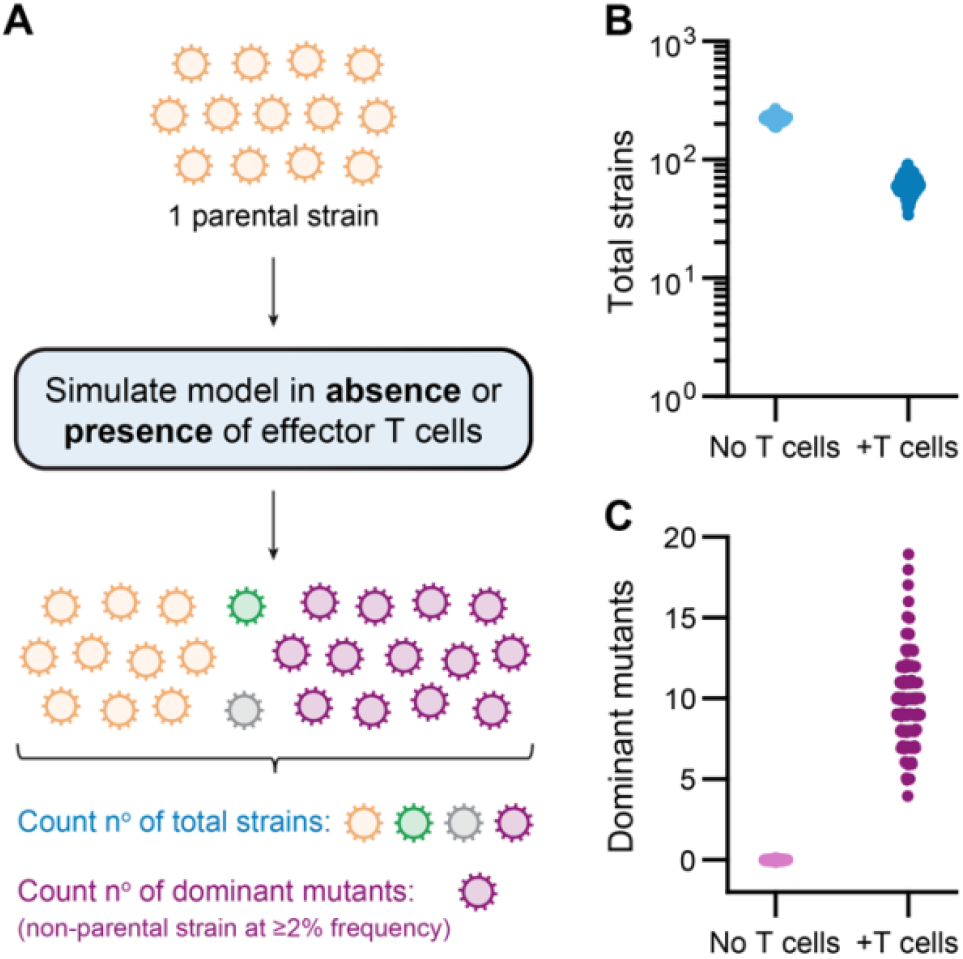
Absence of selective pressure by T cells results in lack of dominant mutants despite high mutation rates. **(A)** Schematic illustrating how model simulations are performed in the presence and absence of effector T cells, starting with a single parental strain. The total (i.e. cumulative) number of strains and the number of dominant mutants (non-parental strains present at a frequency of at least 2% at any point of the simulation) appearing throughout the simulation are quantified. **(B, C)** Total number of strains (B) and number of dominant mutants (C) arising by 200 days post-infection predicted by the model, in the absence of T cells or when the number of T cell clones *N* is set to 100. Data shown result from 200 independent realizations of model simulations.

In light of the apparent discrepancy regarding mutational burden between our model and the aforementioned viral sequence data from RAG2^−/–^ mice (31), we reasoned that a more biologically relevant metric would be the number of viral strains/mutants sufficiently abundant (or “dominant”) relative to the overall pathogen load since these mutants would presumably have a greater impact on the infected host. Furthermore, it is worth noting that variants present at low relative abundances cannot be reliably detected using conventional sequencing techniques. Indeed, it has been estimated that variants present at a frequency below ∼2% are not confidently identified (32), although this threshold value for variant identification is not fixed but rather strongly dependent on the input genome size (33), among other factors. Based on this argument, we instead quantified the number of dominant mutants, defined as the variants of the parental strain that constitute a minimum of 2% of the total pathogen load at any given time throughout the simulation. Indeed, when we quantified the number of dominant mutants predicted by the model with or without effector T cell control, we found that no dominant mutants arose in the absence of T cells, despite a larger number of total strains (**Fig. 3A, C**). This finding thus agrees with previous data showing a lack of measured mutational burden in the absence of selective pressure from T cell-mediated immunity (31) and highlights the importance of distinguishing dominant mutants from other, less abundant variants in subsequent model analyses.

### Intermediate levels of TCR diversity can unexpectedly drive the generation of more dominant mutants that evade the T cell response

Having verified model outcomes at the two ends of the T cell diversity spectrum (i.e., when there are no T cells or there are many unique T cell clones), we next asked how gradually varying the number of T cell clonotypes impacts viral evolution (**Fig. 4A**). In doing so, we found that the total pathogen load and, consequently, the total number of strains that emerge over the course of the simulated 200-day period were inversely proportional to the number of T cell clones in the model (**Fig. 4B, C**), suggesting that more TCR diversity among responding T cells provided increased control of viral replication overall. Interestingly, however, when we quantified the number of dominant mutants as a function of TCR diversity, we found that more mutants emerged on average at intermediate levels of TCR diversity (**Fig. 4D**).

**Figure 4.**
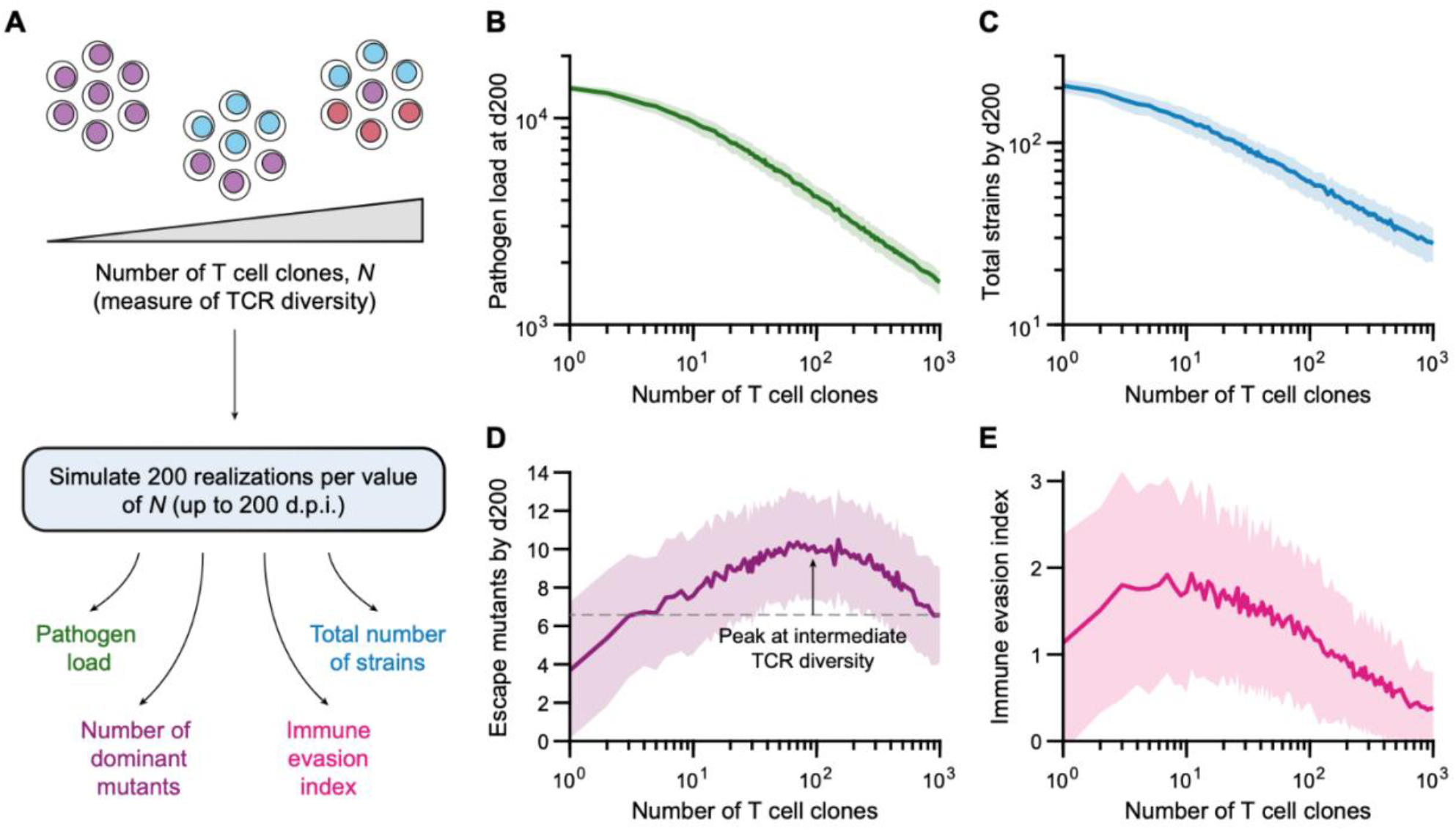
Intermediate level of TCR repertoire diversity in the model promotes greater levels of immune escape. **(A)** Schematic illustrating how model simulations are performed when the number of effector T cell clones, a parameter representing TCR diversity in the model, is gradually varied to test its effect on pathogen replication and evolution over time. **(B, C)** Simulations of pathogen load by day 200 post-infection (B) and total number of pathogen strains that emerges over the course 200 days (C) as a function of the number of T cell clones considered in the model. **(D, E)** Number of dominant mutants (D), defined as the number of strains other than the parental strain with ≥ 2% abundance at any point in time, and pathogen evasion index (E), representing the degree with which mutants evade T cell recognition, as a function of the number of T cell clones. Dashed line in D indicates the average number of dominant mutants for *N* = 100. Curves in panels B-E represent the mean over 200 realizations of model simulations obtained by day 200 post-infection, and shaded areas represent ± standard deviation.

Since we were interested in viral immune escape as a function of TCR diversity, we wondered whether responding T cells in the model were less specific overall for the mutants that do become dominant relative to the parental strain at intermediate TCR diversity values. To investigate this, we developed a metric, the immune evasion index, that measures the degree to which dominant mutants evade recognition by T cells in the model (see Methods). When calculating the value of this index as a function of TCR diversity, we found that intermediate numbers of unique TCR clonotypes not only promoted the formation of more dominant viral mutants (**Fig. 4D**), but that these mutants were less recognizable by responding T cells than dominant mutants emerging at higher TCR diversity (**Fig. 4E**). Importantly, however, this observation was dependent on model parameters; for example, gradually increasing *inter*-clonal competition to be identical to *intra*-clonal competition among responding T cells abolished this effect and the number of dominant mutants instead approached a plateau as TCR diversity was increased (**Fig. S2**). Nonetheless, these data suggest that insufficient TCR repertoire diversification among antigen specific T cells might lead to greater immune escape in certain parameter regimes.

### Replication rate, viral abundance and immune selection pressure all drive the survival of dominant viral mutants

Our observations thus far revealed that mutational burden, as measured by the number of dominant mutants, cannot be predicted simply by quantifying pathogen loads or the total absolute number of mutations. Therefore, we next investigated the reason for the divergent relationship between *dominant* mutant emergence as compared to pathogen load. Based on previous modelling work showing that viral adaptation is maximized when viral abundance is at an intermediate level (15), we hypothesized that an optimal zone of pathogen load exists where the amount of replicating virus is high enough to maintain a high mutation rate, but also low enough that new, more immune evasive mutants can readily overtake prior strains. To directly test this with our model, we checked whether we would observe similar results by gradually increasing equilibrium pathogen loads in the model, while keeping all other parameters constant. Since we do not have an explicit parameter representing pathogen load, this was achieved by varying the pathogen replication rate parameter (*r*_*P*_) (**Fig. 5A**). We found that, whereas pathogen loads and the total number of strains increased monotonically with increasing pathogen replication rates (**Fig. 5B, C**), the number of dominant mutants peaked before falling off sharply when both pathogen replication and loads are high (**Fig. 5D, B**). We observed a similar trend when we quantified the immune evasion index (**Fig. 5E**), which mirrored the number of dominant mutants by peaking at intermediate pathogen replication rates.

**Figure 5.**
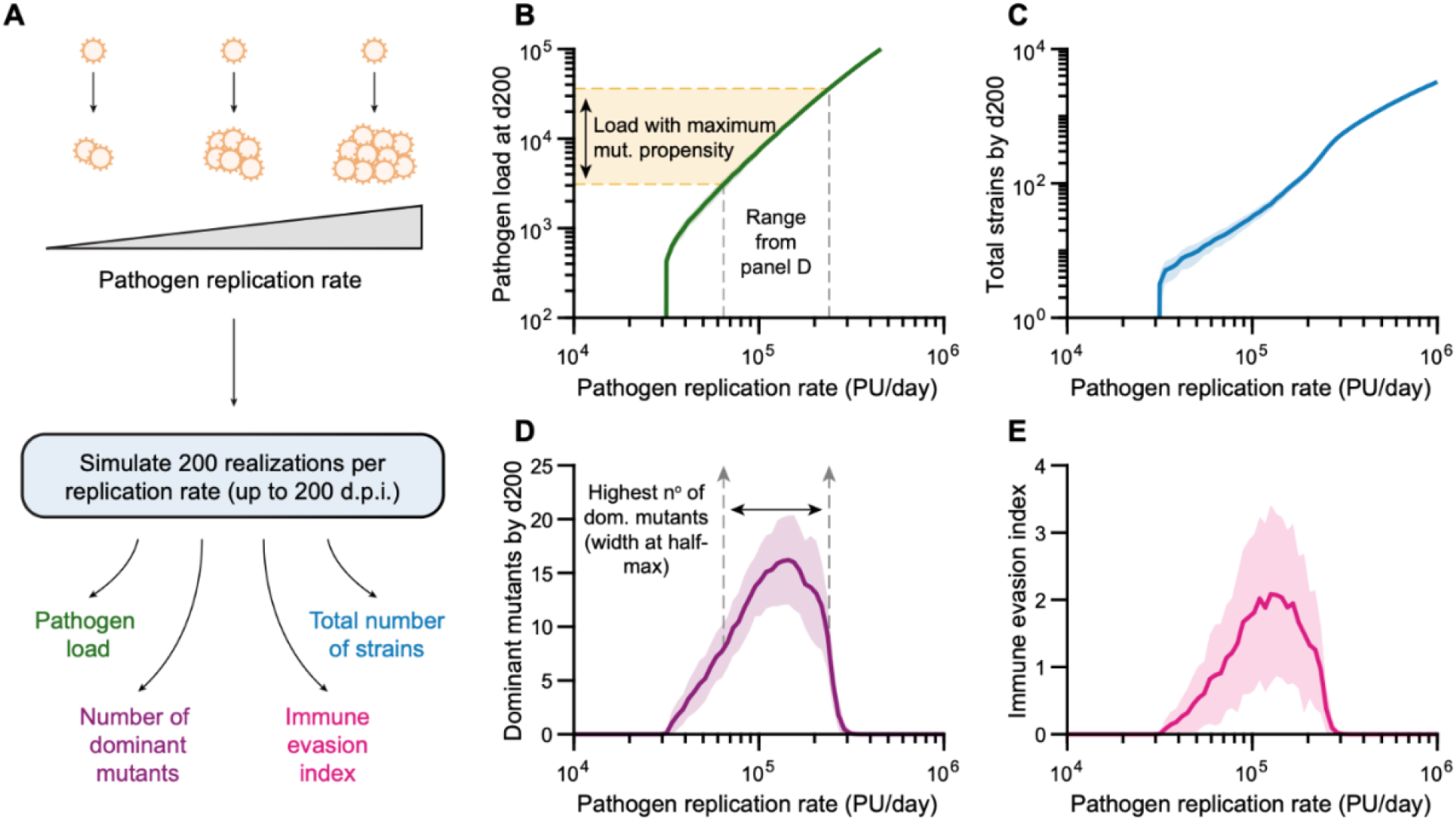
High mutational burden and more immune escape occur within an optimal range of pathogen loads. **(A)** Schematic illustrating how model simulations are performed to investigate the effect of equilibrium pathogen loads on its evolution; the pathogen replication rate (parameter *r*_*P*_ in the model) is gradually increased and model outcomes are assessed as in Fig. 4. The number of T cell clones, *N*, is held constant at a value of 100 clones. **(B, C)** Simulations of pathogen load by day 200 post-infection∼(B) and total number of pathogen strains that emerges over the course of 200 days∼(C) as a function of the rate of pathogen replication. **(D, E)** Number of dominant mutants (D), and pathogen evasion index (E), as a function of pathogen replication rate. The half-maximum width of the curve in D is used to determine the range of pathogen loads in B that maximally promote the emergence of dominant mutants (yellow-shaded region). Curves represent the mean over 200 realizations of model simulations obtained by day 200 post-infection, and shaded areas represent ± standard deviation.

In order to relate the above observations made with respect to the pathogen replication rate parameter *r*_*P*_ back to pathogen loads, we numerically mapped each value of the replication parameter to the resulting pathogen load obtained from model simulations and determined the range of pathogen loads that promotes the highest levels of dominant mutants (**Fig. 5B, D**). We defined this range of pathogen loads (yellow-shaded region in **Fig. 5B**), resulting in more than half of the average maximum number of dominant mutants, as having maximal “mutational propensity”. This pathogen load range exists due to competition for replication between different viral strains of the model, which is a feature of the model that ensures accurate behaviour when multiple strains are present; briefly, when pathogen loads are high, competition for viral replication within target cells makes it less likely for the newly emerging mutants to overtake the existing pool of viral strains. In other words, despite high numbers of mutation events, the relatively lower abundance of newly emerged mutants presents a disadvantage for their replication (i.e., abundance is also a determinant of survival). In summary, the model results suggest that an intermediate range of pathogen loads exists that maximally promotes the emergence of escape mutants, and that this results from a balance between sufficient rates of mutation and competitive dynamics among strains.

### The rate of immune escape depends on a balance between virus-intrinsic dynamics and T cell-mediated immune control

Our results showed that changes in viral dynamics can alter the number of dominant mutants that emerge, with an optimal range of pathogen loads displaying maximal mutational propensity. Next, we asked whether we could tease apart the contributions of the intrinsic properties of the virus versus those that depend on T cell control to immune escape, and how these processes interact to generate the model outcomes observed thus far. To investigate this, we varied the diversity of the T cell response, and for each individual number of T cell clones we repeated the analysis described in the previous section to identify the range of pathogen loads with maximal mutational propensity; doing so allowed us to describe how maximal mutational propensity changes as a function of TCR diversity (**Fig. 6A**). Interestingly, we found that more dominant mutants can emerge at higher numbers of T cell clones within the range of maximal mutational propensity (**Fig. 6B**), representing the greater selection pressure exerted by T cells on viral evolution with increasing TCR diversity.

**Figure 6.**
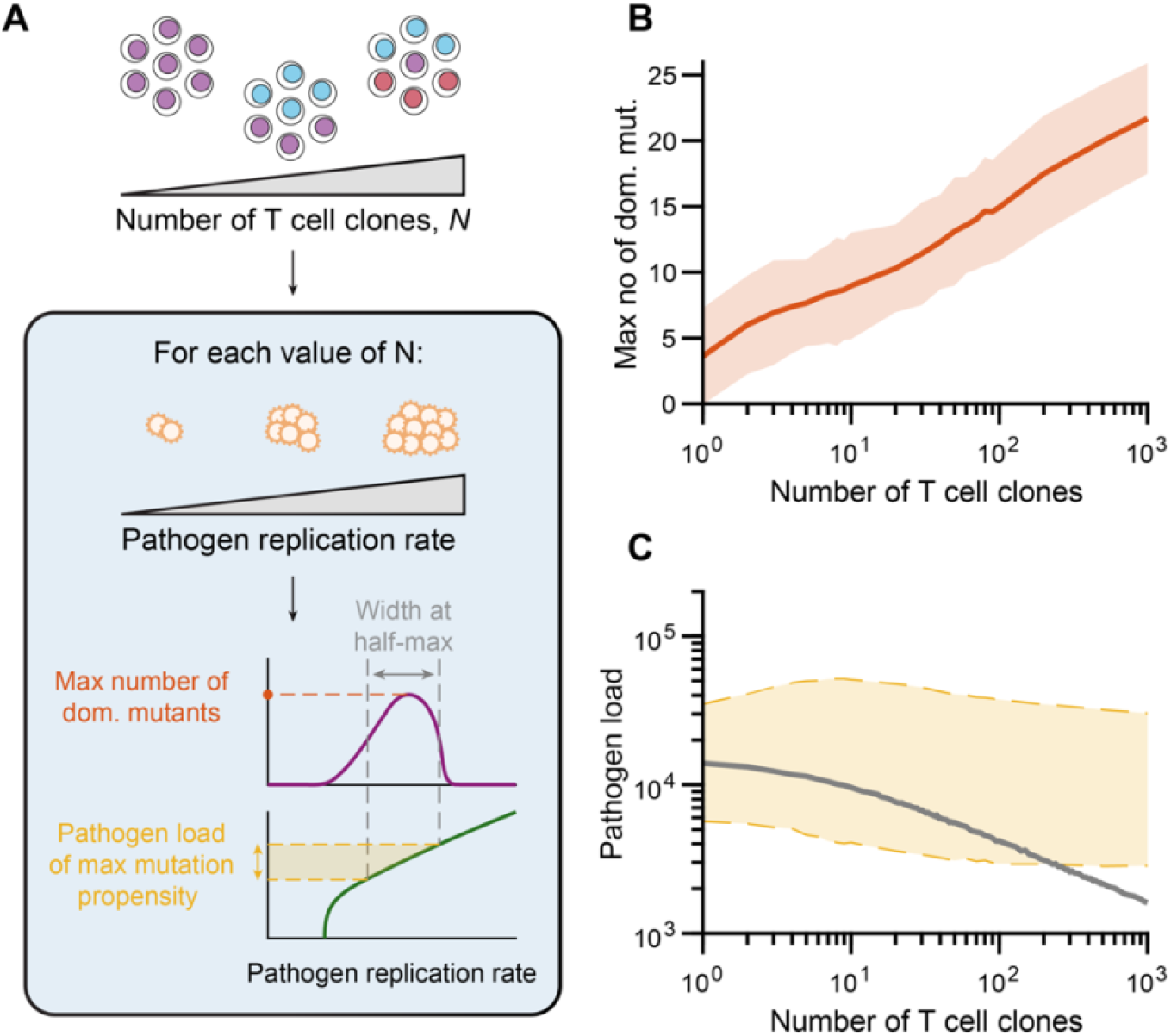
Peak in the number of dominant mutants at intermediate TCR diversities can be explained by the balance between pathogen control and T cell selective pressure. **(A)** A schematic illustrating how model simulations are performed by gradually varying the number of T cell clones, *N*, and at each value of *N* the relationship between pathogen replication rate and the number of dominant mutants is computed as described previously (Fig. 5); the maximum number of dominant mutants and the range of pathogen loads with the highest mutational propensity is determined as a function of *N*. **(B)** The maximum expected number of dominant mutants is higher for higher TCR diversity, indicating that T cell control exerts more selective pressure on pathogens. Curve: mean, shaded region: standard deviation. **(C)** Despite higher selective pressure for large numbers of T cell clones, greater pathogen control decreases computed pathogen abundances (gray curve; identical to Fig. 4B) below the range of pathogen loads with the highest mutational propensity (shaded region bounded by dashed lines).

However, while greater TCR diversity led to more selection pressure on the evolving virus, we previously showed that increasing the number of T cell clones also reduced the total pathogen load (**Fig. 4B**), and that the number of dominant mutants peaked at intermediate TCR diversities before coming back down when the number of unique clones is higher (**Fig. 4D**). We thus wondered whether we could reconcile these findings by comparing the total pathogen load predicted by the model against the range of pathogen loads with maximal mutational propensity, both as a function of TCR diversity. Indeed, by overlaying the observed pathogen load (computed as shown in **Fig. 5B**) onto the computed range within which higher numbers of dominant mutants arise (**Fig. 6C**), we found that increasing the number of T cell clones drove pathogen loads below this range, explaining why fewer dominant mutants are observed than would be expected based purely on selection pressure alone. Of note, we observed that the range of pathogen loads with maximal mutational propensity was relatively independent of TCR diversity, suggesting that this range is largely intrinsic to viral replication dynamics. Taken together, these data suggest that while the optimal range of pathogen loads favouring the emergence of evasive dominant mutants is largely dependent on the intrinsic properties of the virus, different levels of T cell control can alter pathogen loads, and thus mutational burden. Hence, the peak in the rate of immune escape at intermediate TCR diversities can be explained by a balance between higher selective pressure on viral strains and insufficient control of viral replication.

## Discussion

Despite the important role that T cell effector activity plays in shaping viral evolutionary landscapes, it has remained unclear how TCR diversity interacts with virus-intrinsic replication dynamics to promote immune escape. Our modelling results propose that for some ranges of pathogen loads, its propensity for generating mutants that can then efficiently replicate within the host is maximized. We found that, while this range is largely independent of T cell control, it dictates how T cell diversity can influence viral evolution by modulating pathogen loads relative to this range. More specifically, our results showed that while greater T cell control can lead to higher evolutionary selection pressure on the mutating pathogen (driving the potential number of escape mutants up), it also lowers total pathogen loads outside the intrinsic range of maximal mutational propensity and causing the number of escape mutants to go down. Thus, the resulting mutational burden of the virus on the host will depend on the balance between these two opposing forces of T cell mediated control of viral replication and resulting strength of selection. We showed that a potential consequence of these interacting dynamics is a peak in the rate of immune escape at intermediate values of TCR diversity, with fewer dominant mutants on average when the number of responding T cell clonotypes is comparatively lower or higher.

Our observation that the range of pathogen loads with maximal mutational propensity is largely invariant to different levels of T cell selective pressure was intriguing. Earlier theoretical work on viral-immune co-evolution analogously showed that an optimal mutation rate exists that maximizes immune escape, and that, for instance, the experimentally calculated per-genome average mutation rate of HIV-1 lies near this optimum (34). It is thus conceivable that equilibrium pathogen loads of viruses such as HIV have equivalently adapted to lie in a range that maximizes mutational propensity as suggested by our model. Strikingly, however, while the mutation rate per base pair varies by orders of magnitude across different types of DNA and RNA viruses, when accounting for the entire genome size, the per-genome average mutation rate is relatively similar across different species of viruses (35, 36). In spite of this, within-host viral phylogenies still vary enormously (15). While this variation undoubtedly depends on many interconnected aspects of the viral replication cycle within infected cells and the resulting immune response, perhaps viral set points in HIV and HCV, which produce chronic infections in hosts and show substantial evidence of evolution in response to immune pressure (15, 37, 38), fall within these ranges of maximum mutational propensity so as to most successfully evade the host T cell response.

Our results have generated theoretical predictions that could be experimentally tested. For example, mice lacking terminal deoxynucleotidyl transferase (TdT)-dependent TCR repertoire diversification, which possess an estimated 10-fold fewer unique T cell clonotypes (13, 31, 32), could be used to ask whether TdT KO mice are more susceptible to viral immune escape than their wildtype counterparts, as has been previously suggested (39). Alternatively, reconstructed TCR repertoires using retrogenic mice (6, 40) could be used with varying numbers of unique TCRs to ask how increasing the number of unique clonotypes modulates viral control and within-host evolution. Pairing these experimental models with next generation sequencing approaches (32) may enable us to determine how modulation of TCR repertoire diversity affects fitness landscapes and, by extension, the evolutionary trajectories of viruses as they replicate within their hosts. It should be noted, however, that there are significant challenges with these approaches that would first need to be overcome. First, the low frequency and emergence rate of TCR epitope mutations in murine viruses such as LCMV under normal conditions (31) may limit the ability to identify differences generated by varying the number of responding TCR clonotypes. Second, the high stochasticity inherent to the evolutionary process, and observed in our model simulations, would likely require a large number of time and animal resources to be able to resolve how modifying TCR diversity would, on average, affect the rate of within-host viral evolution.

While we kept our model simple in order to focus on the effects of T cell clonal diversity on viral immune escape, other elements that were not included explicitly also exert their effects on the evolutionary dynamics of viruses that are worth noting. For example, our model assumed that pathogen replication rate is unchanged upon mutation, and that there is an equal chance for T cell specificity for the mutant strain to be higher or lower than that of the parent strain. However, it has been established in the context of viral quasispecies evolution that a large number of mutations are deleterious or even lethal for viruses, and that overall fitness tends to decrease across generations (36, 41, 42). Thus, while incorporating the effect of more realistic fitness landscapes would increase the complexity of the model, it certainly remains relevant to ask how incorporating these phenomena into the model could affect predicted outcomes and how TCR diversity profiles impact viral evolution across realistic fitness valleys. Furthermore, it is known that different TCRs may have different propensities for cross-reacting either to slightly altered epitopes with single amino acid mutations (6, 43), or even to broad matrices of unrelated pMHC epitopes as is the case with TCRs that are more germline-like (8). While estimating cross-reactivity on a per-TCR basis is challenging, recent advances has been made in predicting how different TCRs may bind mutant pMHC epitopes (29, 44); expanding on this work in the context of viral infections will be pivotal for estimating the impacts of the number and types of reactive TCRs on the ability for T cells to control the emergence of pMHC epitope mutants and subsequent immune escape.

Our findings could have implications with regard to the generation of escape mutants in chronically infected hosts, which may further impair their ability to contain infection (45, 46). The generation of escape mutants could also have potential epidemiological consequences, providing viruses with the opportunity to seed new hosts with strains that potentially evade pre-existing immunity. For instance, in the context of SARS-CoV-2, it has been proposed that chronically infected individuals have sourced new variants of concern that have hindered containment efforts and spread in spite of high rates of vaccinated and/or previously infected individuals (47, 48). Thus, understanding how different aspects of host immunity, including host TCR repertoire diversity, affects the likelihood of the generation of these mutants is an important step in predicting which factors maximally drive viral evolution both within and between hosts. In summary, by applying the principles of viral-immune co-evolution to specifically study the impact of TCR diversity on immune escape, interesting predictions arose from our model results that suggest TCR repertoire diversity can modulate the propensity for viruses to evade the host response independently of mutation rates (which were held constant in our model simulations). Our observations led to intriguing insights that may have broader implications for the drivers of not only within-host evolution but also potentially for viral adaptations occurring at epidemiological scales.

## Methods

### Model formalism

To model the dynamics of pathogen evolution as a function of TCR repertoire diversity, we constructed an ordinary differential equation (ODE) model consisting of *N* fixed clones of effector T cells and *M*(*t*) strains of a mutating pathogen *P*_*j*_(*t*) (with *j* ∈ (1, 2, …, *M*(*t*))), where the total number of pathogen strains, *M*(*t*), is allowed to increase with time via random mutation events. The system of equations describing the model can be written as

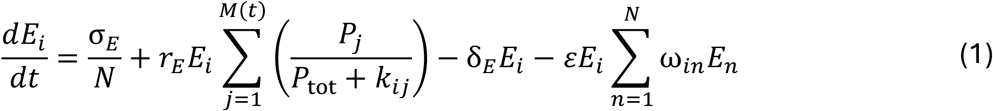

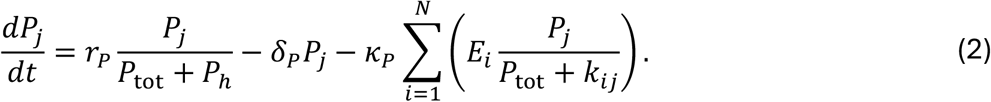

Equation (1), describing the rate of change of effector T cell clone *i* over time, was adapted from previous models of T cell dynamics (49-52). The source term σ_*E*_ represents total thymic input of all pathogen-specific T cells; although different clonotypes are present at different precursor frequencies in the naïve T cell repertoire (7), for simplicity we let thymic input be equally divided among all *N* T cell clones. All clones are assumed to share a natural turnover rate given by δ_*E*_, and to proliferate with a maximum rate *r*_*E*_ upon pathogen recognition. The contribution of each strain of pathogen, *P*_*j*_, will depend on the abundance of that strain relative to the total pathogen load, denoted by *P*_tot_ = ∑_*j*_ *P*_*j*_, and on the specificity of T cell clone *i* for strain *j*, a value that depends reciprocally on the magnitude of the parameter *k*_*ij*_. This latter parameter is sampled randomly from a log-normal distribution, with base-10 logarithmic mean μ_*k*_ and standard deviation σ_*k*_. The final term in Eq. (1) represents the rate of intra- and inter-clonal competition among T cells. We set the intra-clonal competition rate to ε, which represents competition for cognate antigen as well as other survival signals (49, 50, 53), and let inter-clonal competition occur at a lower rate *f*ε, where 0 < *f* < 1 (since different T cell clones may have different antigen specificity). In other words, we assumed that

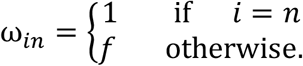

We described the rate of change of each pathogen strain *P*_*j*_ as a function of time by Eq. (2). Specifically, we assumed that the *per capita* replication rate of each strain is bounded above by the fraction *r*_*P*_/*P*_*h*_, and that, when pathogen loads are sufficiently high (i.e., when *P*_tot_ > *P*_*h*_), more abundant strains have a higher probability of infecting target cells and thus replicate more efficiently. We let δ_*P*_ be the rate that encompasses all T cell independent mechanisms of pathogen clearance, while the final term in Eq. (2) represents T cell-mediated pathogen clearance of strain *j* at a maximum rate of κ_*P*_ per T cell clone *i*. We assumed this rate to be dependent again on the specificity of clone *i* for strain *j*, represented by the reciprocal of *k*_*ij*_, as well as the relative abundance of strain *j*.

### Mutation events

Generation of mutant strains in the model was obtained using a non-homogeneous Poisson point process, where the stochastic frequency of novel variant formation from a parental strain was assumed to be proportional to the abundance of that strain, modulated by an average *per capita* mutation rate. To do this, at each time step Δ*t* in the simulation, we sampled a number *M*_new_ arising from strain *P*_*j*_ from a Poisson distribution with a rate parameter λ(*t*) = *r*_*mut*_ *P*_*j*_(*t*) Δ*t*, similar to previous modelling work on HIV evolution (45). To prevent long computational times, we did not allow new variants to be produced once the total number of strains exceeded a high number *M*_max_.

When a new variant, *P*_*m*_, is formed from *P*_*j*_, we assumed that the specificity of each T cell clone *i* for the new strain *m* is perturbed relative to that of clone *i* for the parental strain *j*. To implement this, we let the magnitude of the perturbation be drawn from a log-normal distribution, with a standard deviation θ_*k*_ and centered at 0 in base-10 logarithmic space, i.e., log_10_(*k*_*im*_) = log_10_(*k*_*ij*_) + 𝒩(0, θ_*k*_). This is illustrated schematically in Fig. 1C, where the specificity of each clone *i* for a given strain was assessed via the reciprocal of the parameter *k*_*ij*_ or *k*_*im*_ (for the parental strain *j* or the mutant strain *m*, respectively).

### Immune evasion index

The immune evasion index conceptually represents the degree of pathogen evolution and is modulated both by the number of dominant mutants and by the reduced ability for T cells to recognize those mutants relative to the parental (or originally infecting) strain of the pathogen. Let 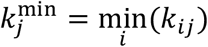 represent the reciprocal of the highest specificity of any effector T cell clone for pathogen strain *j* (lower values of *k*_*ij*_ indicate higher specificity of clone *i* for strain *j*, thus the lowest value of *k*_*ij*_ for a given *j* represents the highest specificity across all clones for strain *j*). For the parental strain, this is equivalent to 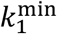. The immune evasion index (IEI) is then defined by

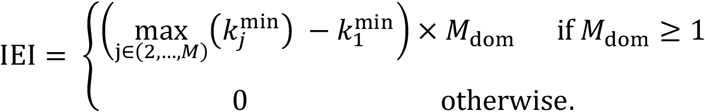

Here, *M*_dom_ represents the number of dominant mutants.

### Model parameters and numerical implementation

**Table S1** summarizes the different parameters of the model and their respective values used in the simulations. These parameters and their values are similar to our previous work analyzing the dynamics of effector T cells responding to replicating pathogen (50) using a computational model that was fit to serum viral load data from LCMV-Cl13 infected mice (54).

In order to remove transient artifacts arising from improper initialization of the model, we first simulated the model in the absence of pathogen to allow the effector T cell population to reach a steady state; these steady state values were then used as the initial condition for effector T cells upon introduction of the parental strain of the pathogen in the model, where we set *P*_1_(*t* = 0) → 1 PU (pathogen unit). Similarly, when a new strain *m* is produced via a mutation event at time *t*_*m*_, we set *P*_*m*_(*t* = *t*_*m*_) → 1 PU. MATLAB version 2022a was used to write all codes and generate all numerical results pertaining to the model.

### LCMV-Cl13 infections in mice

C57BL/6 and C57BL/6 TCRβ^−/–^ mice (55) were purchased from Jackson Laboratories (Bar Harbor, ME), and then bred in-house. Infections were performed at 6-8 weeks of age with only females. Animal housing, care, and research were in accordance with the Guide for the Care and Use of Laboratory Animals and all procedures performed were approved by the McGill University Animal Care Committee (protocol #MCGL-7570). The Clone 13 strain of LCMV was propagated from stocks provided by Martin Richer (University of Indiana) on L929 cells, as described previously (50). Mice were infected by injecting 2 × 10^6^ plaque forming units (PFU) of LCMV-Cl13 intravenously as per established protocol (54, 56). Mice were euthanized 21 days post-infection, and their spleens were collected in 1% RPMI before being weighed and homogenized in Lysing Matrix D tubes (MP Biomedicals) using a MagNA Lyser (Roche) at 6000 rpm for 40 seconds. Spleen homogenate was then spun down at 12,000 rpm for 10 minutes at 4 °C. To further clarify the solution, the supernatant was transferred to a new sterile tube and re-spun at 12,000 rpm for 10 minutes at 4 °C, and 400 μL of TRIzol reagent (Invitrogen) was added to 200 μL aliquots of the supernatant from homogenized spleens and stored at –80 °C for RNA processing and quantification of viral titres. To quantify viral loads in the spleens of WT or TCRβ^−/–^ mice, samples were thawed on ice, 80 μL of chloroform added, then incubated at room temperature for 5 minutes, and spun down at 12,000 rpm for 15 minutes. RNA was extracted using the PureLink RNA Mini Kit (Invitrogen), followed by cDNA conversion of 220 ng of RNA in 20 μL reaction volumes with the High-Capacity cDNA Reverse Transcription Kit (Invitrogen). Viral titres were determined via qPCR to quantify LCMV-GP RNA levels using primers described previously (57).

## Acknowledgements

We would like to thank the animal facility staff at McGill University for their excellent care of our animal colony. H.J. was supported by an Alexander Graham Bell Canada Graduate Scholarship - Doctoral (NSERC) and a B2X Doctoral Award (FRQNT). J.N.M. is a Canada Research Chair for Immune Cell Dynamics. This work was supported by an NSERC Discovery Grant #2019-04520 (A.K.) and a McGill Mi4 Round 5 seed grant (J.N.M.).

