## Supplemental figures and text for "Within-host viral evolution varies with T cell receptor repertoire diversity independently of viral mutation rates"

### Supplementary Figures and Tables

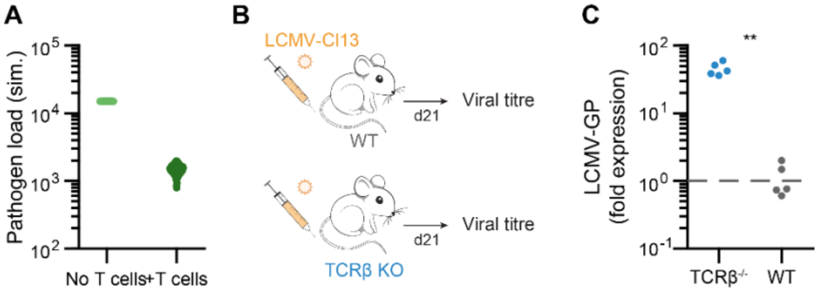


**Figure S1.** Lack of host T cells leads to higher pathogen loads both *in silico* and *in vivo*. **(A)** Pathogen loads at day 15 post-infection in the model when simulated in the absence or presence of effector T cells. **(B)** WT and TCRβ KO mice (n = 5 each) were infected with LCMV-Cl13, and spleens were harvested at day 14 for viral titre determination. **(C)** LCMV-GP RNA relative fold expression, assessed by qPCR. *P*-value computed using non-parametric Wilcoxon rank-sum test.

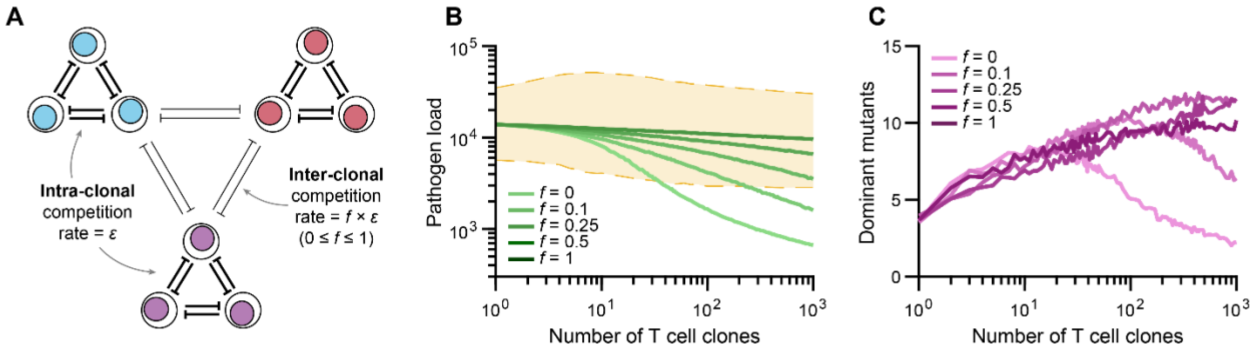


**Figure S2.** Symmetrical competition within and between T cell clones abolishes peak in mutational burden at intermediate TCR diversity. **(A)** Schematic depicting intra- vs. inter-clonal T cell competition. The inter-clonal competition is assumed to be a fraction, $0 \leq f \leq 1$, of the inter-clonal competition rate $\varepsilon$. **(B)** Increasing $f$ from 0 (no inter-clonal competition) to 1 (symmetrical competition) reduces the effect of the number of clones $N$ on pathogen control. **(C)** The number of dominant mutants does not peak as a function of TCR diversity when intra- and inter-clonal competition are similar, i.e., as $f \rightarrow 1$.

**Table S1.** Parameter values used in model simulations. These values were chosen to be similar to those in our previous modeling work of T cell control of replicating pathogen (Ref. 50 in main text). PU = pathogen units.

| Param | Description | Value(s) | Units |
|---|---|---|---|
| $\sigma_E$ | Thymic input across all TCR clonotypes | 150 | cells day$^{-1}$ |
| $r_E$ | Maximum proliferation rate of effector T cells | 4.0 | day$^{-1}$ |
| $k_{ij}$ | Pathogen load of strain j evoking half-maximum response of T cell clone i, drawn from a log-normal distribution | Range | PU$^{-1}$ |
| $\mu_k$ | Base-10 logarithmic mean used to sample $k_{ij}$ | 4.3 | – |
| $\sigma_E$ | Base-10 logarithmic standard deviation used to sample $k_{ij}$ | 0.25 | – |
| $\delta_E$ | Natural turnover rate of effector T cells | 0.3 | day$^{-1}$ |
| $\varepsilon$ | Intra-clonal competition rate | 3.2 x $10^{-6}$ | cells day$^{-1}$ |
| $f$ | Scaling factor for inter-clonal competition | 0.1 | – |
| $N$ | Number of unique pathogen-specific TCR clonotypes | Range | – |
| $M_0$ | Initial number of pathogen strains | 1 | – |
| $r_P$ | Maximum replication rate | 7.5 x $10^4$ | PU day$^{-1}$ |
| $P_h$ | Half-maximum constant of pathogen replication | 1 x $10^4$ | PU |
| $\delta_P$ | T cell-independent pathogen clearance rate | 3 | PU day$^{-1}$ |
| $\kappa_P$ | Maximum pathogen removal rate per T cell | 0.015 | PU (cell day)$^{-1}$ |
| $r_{mut}$ | Per capita average mutation rate used to randomly generate new pathogen strains | 7.5 x $10^{-5}$ | day$^{-1}$ |
| $\theta_k$ | Base-10 logarithmic standard deviation used to sample perturbations of $k_{ij}$ from parent to daughter strain | 0.1 | – |
| $M_{max}$ | Maximum allowable number of mutant strains per simulation | $10^4$ | – |

**Supplementary Text**

The system of ODEs that describes the computational model in Eqs. (1)-(2) in the main text needed to satisfy two important limits. The first limit considers the case where all effector T cell clones are identical and therefore have the same specificity for a given strain $j$. That is,

$$k_{1j} = k_{2j} = \cdots = k_{Nj} \quad \equiv \quad k_{*j}.$$

In this case, the competition between these identical T cell clones would be fully symmetrical (since they would have the same antigen specificity), i.e., $\omega_{in} = 1,\ \forall\, i, n \in \{1, \dots, N\}$. Within this framework, the model should also be equivalent to a model with a single effector T cell clone ($N = 1$), denoted by $\tilde{E}$. To check this, we verified that the dynamics of the sum of identical clones, $E_{tot} = \sum_i E_i$, are the same as those produced by $\tilde{E}$ whose specificity for strain $j$ is $1/k_{*j}$. The dynamics of this latter one-clone model are given by

$$\frac{\mathrm{d}\tilde{E}}{\mathrm{d}t} = \sigma_E + r_E \tilde{E} \sum_{j=1}^{M(t)} \left( \frac{P_j}{P_{\mathrm{tot}} + k_{*j}} \right) - \delta_E \tilde{E} - \varepsilon \tilde{E}^2 \tag{S1}$$

$$\frac{\mathrm{d}P_j}{\mathrm{d}t} = r_P \frac{P_j}{P_{\mathrm{tot}} + P_h} - \delta_P P_j - \kappa_P \tilde{E} \frac{P_j}{P_{\mathrm{tot}} + k_{*j}}. \tag{S2}$$

By substituting $k_{ij} = k_{*j}$ and $\omega_{in} = 1$ in the original model of Eq. (1) and writing it with respect to $\mathrm{d}E_{\mathrm{tot}}/\mathrm{d}t$, we obtain

$$\begin{aligned} \frac{\mathrm{d}E_{\mathrm{tot}}}{\mathrm{d}t} &= \sum_{i=1}^{N} \frac{dE_i}{dt} \\ &= \sum_{i=1}^{N} \left( \frac{\sigma_E}{N} + r_E E_i \sum_{j=1}^{M(t)} \left( \frac{P_j}{P_{\mathrm{tot}} + k_{*j}} \right) - \delta_E E_i - \varepsilon E_i \sum_{n=1}^{N} E_n \right) \\ &= \sigma_E + r_E E_{\mathrm{tot}} \sum_{j=1}^{M(t)} \left( \frac{P_j}{P_{\mathrm{tot}} + k_{*j}} \right) - \delta_E E_{\mathrm{tot}} - \varepsilon E_{\mathrm{tot}}^2, \end{aligned}$$

which, when setting $E_{tot} = \tilde{E}$, matches exactly with Eq. (S1). Similarly, by applying these substitutions to Eq. (2), we obtain

$$\begin{aligned} \frac{\mathrm{d}P_j}{\mathrm{d}t} &= r_P \frac{P_j}{P_{\mathrm{tot}} + P_h} - \delta_P P_j - \kappa_P \sum_{i=1}^{N} \left( E_i \frac{P_j}{P_{\mathrm{tot}} + k_{*j}} \right) \\ &= r_P \frac{P_j}{P_{\mathrm{tot}} + P_h} - \delta_P P_j - \kappa_P E_{\mathrm{tot}} \frac{P_j}{P_{\mathrm{tot}} + k_{*j}}. \end{aligned}$$

This in turn matches exactly with Eq. (S2) when setting $E_{\text{tot}} = \tilde{E}$. It follows that when all effector T cell clones are identical, the two models are equivalent, and the first limiting case is satisfied.

In the second limiting case, we apply a similar logic by making all pathogen strains to be identical, implying that any given T cell clone $i$ will be equally specific for all strains. By ignoring the time dependency of the maximum number of strains $M$ resulting from any new mutation event, we have

$$k_{i1} = k_{i2} = \cdots \quad \equiv \quad k_{i*}.$$

In this case, the model should be equivalent to a model with a single pathogen strain ($M = 1$), denoted by $\tilde{P}$. As before, we verified that the dynamics of the sum across all strains, $P_{tot} = \sum_j P_j$, are the same as those produced by $\tilde{P}$, where the specificity of T cell clone $i$ for $\tilde{P}$ is equal to $1/k_{i*}$, by considering the one-strain model given by

$$\frac{dE_i}{\mathrm{d}t} = \frac{\sigma_E}{N} + r_E E_i \frac{\tilde{P}}{\tilde{P} + k_{i*}} - \delta_E E_i - \varepsilon E_i \sum_{n=1}^{N} \omega_{in} E_n, \tag{S3}$$

$$\frac{\mathrm{d}\tilde{P}}{\mathrm{d}t} = r_P \frac{\tilde{P}}{\tilde{P} + P_h} - \delta_P \tilde{P} - \kappa_P \sum_{i=1}^{N} \left( E_i \frac{\tilde{P}}{\tilde{P} + k_{i*}} \right). \tag{S4}$$

By substituting $k_{ij} = k_{i*}$ in Eq. (1) of the original model and ignoring the time dependency of $M$, we obtain

$$\begin{aligned}\frac{dE_i}{dt} &= \frac{\sigma_E}{N} + r_E E_i \sum_{j=1}^{M} \left( \frac{P_j}{P_{\text{tot}} + k_{i*}} \right) - \delta_E E_i - \varepsilon E_i \sum_{n=1}^{N} \omega_{in} E_n \\ &= \frac{\sigma_E}{N} + r_E E_i \frac{P_{\text{tot}}}{P_{\text{tot}} + k_{i*}} - \delta_E E_i - \varepsilon E_i \sum_{n=1}^{N} \omega_{in} E_n,\end{aligned}$$

which, when setting $P_{\text{tot}} = \tilde{P}$, matches exactly Eq. (S3). Similarly applying these substitutions to Eq. (2) and writing it with respect to $dP_{\text{tot}}/dt$, we obtain

$$\begin{aligned}\frac{dP_{tot}}{dt} &= \sum_{j=1}^{M} \frac{dP_j}{dt} \\ &= \sum_{j=1}^{M} \left( r_P \frac{P_j}{P_{\text{tot}} + P_h} - \delta_P P_j - \kappa_P \sum_{i=1}^{N} \left( E_i \frac{P_j}{P_{\text{tot}} + k_{i*}} \right) \right) \\ &= r_P \frac{P_{\text{tot}}}{P_{\text{tot}} + P_h} - \delta_P P_{\text{tot}} - \kappa_P \sum_{i=1}^{N} \left( E_i \frac{P_{\text{tot}}}{P_{\text{tot}} + k_{i*}} \right).\end{aligned}$$

This in turn matches exactly Eq. (S4) when setting $P_{tot} = \tilde{P}$. This means that when all strains are identical, the two models are equivalent, and the second limiting case is also satisfied.

Note that the second limiting case would not have been satisfied if $P_{\text{tot}}$ in the denominator of the terms involved in effector T cell proliferation and T cell-dependent pathogen clearance (i.e., in $P_{tot} + k_{ij}$), were replaced with $P_j$. In other words, if the proliferation of, and clearance by, T cells that are induced by a single pathogen strain were completely independent from the presence and abundance of other variants of that pathogen, then the model would depend arbitrarily on the number of strains despite all variants being physiologically indistinguishable. Similarly, if we had replaced $P_{tot}$ in the denominator of the replication term of pathogen dynamics with $P_j$, the second limiting case would also no longer be satisfied.

By accounting for these limits when developing the model, we avoided generating results that were an artifact of its design. This form introduces competition between different strains of the pathogen since, for example, the replication of a lowly abundant variant is disfavoured over a prominent one as a result of $P_{tot}$ in the denominator of pathogen replication in Eq. (2); this in turn gives rise to an interesting result of the model, which is the pathogen intrinsic range of maximum mutational propensity (Figs. 5 and 6). This is because, when the pathogen load of an abundant strain is too high, its competitive advantage makes it less likely for other variants to overtake it and thus for additional dominant mutants to arise. If, on the other hand, pathogen replication of a given strain were completely independent from the presence and abundance of other strains, we would expect the number of dominant mutants to more closely resemble the total number of strains arising from the simulations, i.e., we would no longer see a mismatch between these two metrics as we did in our model (Fig. 3). This reasoning provides support for our model prediction that T cell-dependent mechanisms contribute to, but do not fully account for, immune escape, and that these work together with pathogen intrinsic properties to modulate the abundance and evasiveness of escape mutants.
